# NMDA-dependent coplasticity in VIP interneuron-driven inhibitory circuits

**DOI:** 10.1101/2025.01.07.631664

**Authors:** Jadwiga Jabłońska, Grzegorz Wiera, Jerzy W. Mozrzymas

**Affiliations:** Department of Biophysics and Neuroscience, Wroclaw Medical University, 3a Chalubinskiego Str., 50-368 Wroclaw, Poland

**Keywords:** GABA, iLTP, inhibitory plasticity, VIP, somatostatin, parvalbumin, inhibitory synapse, stratum oriens, NMDA, coplasticity, plasticitome

## Abstract

Inhibitory plasticity is emerging as a key regulator of excitation/inhibition (E/I) balance, a fundamental determinant of brain network dynamics. While significant progress has been made in understanding inhibitory plasticity at synapses targeting excitatory principal neurons (I→E), the mechanisms and functional implications of plasticity at interneuron-interneuron (I→I) synapses remain largely unexplored. Herein, we investigated the properties and plasticity of inhibitory inputs from vasoactive intestinal peptide (VIP) interneurons onto *stratum oriens* interneurons (*so*INs) in the hippocampal CA1 region. Using optogenetics, patch-clamp electrophysiology, and morphological reconstructions, we characterized the kinetics, short-term plasticity, and NMDA receptor-dependent long-term plasticity at VIP→*so*IN synapses in two distinct *so*IN subtypes: fast-spiking (FS) and oriens-lacunosum moleculare (OLM)/bistratified interneurons. Optogenetically evoked VIP→*so*IN IPSCs showed faster rise times and slower decay kinetics in FS interneurons compared to OLM/bistratified cells, although both subtypes exhibited similar short-term plasticity profiles. Brief NMDA receptor activation (1 min) induced long-term depression (iLTD) at VIP→OLM/bistratified synapses, but not at VIP→FS synapses, underscoring subtype-specific plasticity. However, prolonged NMDA exposure (2 min) elicited iLTD in both interneuron subtypes. Interestingly, excitatory inputs to *so*INs demonstrated NMDA-induced long-term potentiation (E→I LTP) after brief NMDA exposure, but not after prolonged application. Notably, coplasticity analysis in individual *so*INs revealed asymmetric co-expression of I→I LTD and E→I LTP in OLM/bistratified interneurons. In contrast, FS interneurons exhibited a duration-dependent transition between asymmetric and symmetric coplasticity. These findings reveal a target-cell-specific landscape of inhibitory I→I plasticity and its co-expression with excitatory plasticity, highlighting VIP interneurons as key modulators of E/I balance within local hippocampal circuits.

## Introduction

The brain’s ability to perform diverse cognitive functions relies fundamentally on neuroplasticity, encompassing various types of plastic changes, including structural rearrangements of neuronal networks, alterations in endogenous excitability, and synaptic plasticity. The latter process critically regulates the subtle balance between excitation and inhibition, a key feature shaping the outcome of neuronal operations. Although much research has historically focused on excitatory synapses, the critical role of inhibitory synaptic plasticity in maintaining excitation-inhibition (E/I) balance, stabilizing neural networks, and driving circuit plasticity is increasingly being recognized (Hennequin et al., 2017; Wu et al., 2022). One particularly intriguing aspect of inhibitory plasticity is its reliance on heterosynaptic mechanisms, where changes at inhibitory synapses depend on concurrent plasticity at neighboring synapses, often excitatory in nature (Barberis, 2020; Ravasenga et al., 2022; McFarlan et al., 2023; Wiera et al., 2024). This interdependence, referred to as coplasticity, allows inhibitory plasticity to be integrated into broader network activity, enabling precise coordination and fine-tuning of neuronal responses.

Studies of the mechanisms of inhibitory plasticity are inherently challenging, and the diversity of GABAergic synapses is one of the main complicating factors. This diversity stems from the numerous types of inhibitory interneurons, forming synapses with distinct functional properties and targeting specific neuronal compartments of both excitatory and inhibitory neurons (Pelkey et al., 2017, for the hippocampus). In recent years, significant progress has been made in elucidating the mechanisms of plasticity at inhibitory synapses onto excitatory pyramidal neurons, commonly referred to as I→E synapses (Petrini et al., 2014; Chiu et al., 2018; Field et al., 2020; Wiera et al., 2024). However, far less is known about plastic changes at inhibitory synapses that target other inhibitory neurons (I→I) despite their critical role in shaping neural circuit activity (Karnani et al., 2016). Interneuron-interneuron synapses have been recognized as key modulators of network dynamics, particularly through disinhibitory pathways that transiently increase excitatory activity (Kullander and Topolnik, 2021; Ferguson et al., 2023; Veit et al., 2023). Vasoactive intestinal peptide (VIP)-positive interneurons stand out as a specialized class of GABAergic cells with predominantly disinhibitory functions. These interneurons selectively inhibit other inhibitory cells, such as *oriens-lacunosum moleculare* (OLM) interneurons and basket cells of the hippocampal CA1 field. This I→I→E connectivity motif enables dynamic circuit-level flexibility by gating excitation on principal neurons (Pfeffer, 2014; Tyan et al., 2014).

Alongside their emerging disinhibitory role, VIP interneurons are also crucial for shaping higher-order cognitive functions, such as attention, learning, memory, and brain rhythms (Fu et al., 2014; Adler et al., 2019). Despite their central role in the regulation of hippocampal and cortical circuits, little is known about the plasticity of VIP-mediated inhibition. Putative plasticity at VIP I→I synapses is particularly interesting given the diversity of their postsynaptic targets, which raises the possibility of synapse-specific plasticity rules. For example, in the motor cortex, the causal spike timing-dependent protocol (STDP) induces long-term depression (LTD) at VIP→Martinotti cell synapses, whereas VIP→basket cell synapses do not exhibit detectable plasticity in response to the same stimulation (McFarlan et al., 2024b). This synapse-specific I→I plasticity may support the differential roles of distinct VIP synapses in modulating local inhibitory circuits (Magnin et al., 2019; Kullander and Topolnik, 2021). Similarly, inhibitory interneurons show a wide range of plasticity mechanisms at their excitatory inputs (Chistiakova et al., 2019; Lamsa and Lau, 2019; Honoré et al., 2021). However, the interplay, or coplasticity, between VIP-mediated inhibitory and excitatory synaptic inputs remains unexplored.

NMDA receptors are widely recognized as essential players in shaping both excitatory and inhibitory synaptic plasticity in the hippocampus (Muir et al., 2010; Chiu et al., 2018; Wiera et al., 2021; Brzdąk et al., 2023). Their activation sets off a cascade of events driven by calcium influx, which orchestrates key processes, such as CaMKII or calcineurin activation and the rearrangement of gephyrin, which critically regulate long-term changes in the strength of GABAergic synapses (Petrini et al., 2014; Wiera et al., 2022). Notably, NMDA-induced plasticity is particularly relevant to the concept of intersynaptic coplasticity, where simultaneous changes in inhibitory and excitatory inputs fine-tune the excitatory/inhibitory (E/I) balance both within individual cells and across neural circuits (Ravasenga et al., 2022; Wiera et al., 2024). While much has been documented regarding input-specific NMDA-dependent coplasticity in pyramidal cells, its presence at interneuron synapses remains relatively unexplored. To address this gap, we focused on NMDA-induced plasticity at VIP-positive synapses targeting CA1 *stratum oriens* interneurons (*so*INs). Using a combination of optogenetic stimulation, patch-clamp electrophysiology, and morphological reconstructions, we investigated the I→I plasticity profiles of two key interneuron subtypes: fast-spiking (FS) interneurons and OLM/bistratified cells. Additionally, we examined the co-expression of plasticity at VIP inhibitory inputs and excitatory synapses following different durations of NMDA administration. Our results show that NMDA treatment induced both inhibitory (VIP IN input) and excitatory plasticity in synapses located on analyzed *so*INs, with these effects strongly dependent on the timing of NMDA exposure. This reveals a potential mechanism for regulating E/I balance and provides new insights into how intersynaptic coplasticity contributes to the dynamic remodeling of hippocampal circuits (McFarlan et al., 2023). Finally, by characterizing I→I-specific plasticity, this study advances our understanding of VIP-driven modulation in hippocampal circuits and may shed new light on phenomena of network adaptation and cognitive processes.

## Materials and methods

### Animals

VIP-Cre::Ai32 mice expressing channelrodopsin-2 in VIP-positive neurons were generated by crossing parental lines Ai32 (B6.Cg-Gt(ROSA)26Sortm32(CAG-COP4*H134R/EYFP)Hze/J; Jackson Labs stock #024109) and VIP-IRES-Cre (B6J.Cg-Viptm1(cre)Zjh/AreckJ; Jackson Labs stock #031628). Animals were housed under a nonreversed 12h/12h light-dark cycle with water and food supplied ad libitum. Both male and female animals aged P60 to P150 were used in the experiments. All animal procedures in this study complied with Polish regulations, specifically the Act on the Protection of Animals Used for Scientific or Educational Purposes (Act of January 15, 2015, with subsequent amendments), as well as the EU Directive 2010/63/EU. Experiments involving genetically modified organisms were authorized by the Polish Ministry of Environment under approval numbers 144/2018 and 69/2023.

### Slice preparation

Acute brain slices were prepared as previously described (Jabłońska et al., 2024; Wiera et al., 2024). Briefly, mice were anesthetized using isoflurane and subsequently decapitated. The brains were promptly extracted and immersed in ice-cold, oxygenated (95 % O₂/5 % CO₂) holding artificial cerebrospinal fluid (holding aCSF) containing (in mM): 92 NaCl, 2.5 KCl, 1.2 NaH₂PO₄, 30 NaHCO₃, 20 HEPES, 25 D-glucose, 5 sodium ascorbate, 2 thiourea, 3 sodium pyruvate, 1.3 MgSO₄, and 2.5 CaCl₂, pH 7.4. Slices (350 µm-thick) were sectioned horizontally using a vibrating blade-microtome (VT1200S, Leica) and transferred to a recovery chamber containing NMDG-based recovery solution (in mM: 93 N-methyl-D-glucamine, 2.5 KCl, 1.2 NaH₂PO₄, 30 NaHCO₃, 20 HEPES, 25 glucose, 5 sodium ascorbate, 2 thiourea, 3 sodium pyruvate, 10 MgSO₄, 0.5 CaCl₂, pH 7.4) maintained at 34 °C. During a 30-minute recovery period, increasing volumes (0.25-2 mL) of 2 M NaCl were added to the recovery chamber to gradually adjust the concentration of Na^+^ ions (Ting et al., 2018). Slices were subsequently transferred to holding aCSF and maintained at room temperature until use. Electrophysiological experiments were conducted within 8 hours of slice preparation.

### Electrophysiological recordings

At least 1 h after preparation, hippocampal slices were transferred to a submerged recording chamber and continuously perfused at 2–3 mL/min with oxygenated aCSF (in mM: 119 NaCl, 26.3 NaHCO₃, 11 glucose, 2.5 KCl, 1 NaH₂PO₄, 1.3 MgSO₄, 2.5 CaCl₂, pH 7.4, 295−300 mOsm), and the temperature was maintained at 28 °C. The health, and cell body shape of the targeted interneurons in the CA1 *stratum oriens* were assessed using a Zeiss Axio Examiner Z1 microscope with infrared differential interference contrast and a 40x objective. Whole-cell patch-clamp recordings were performed with borosilicate glass capillaries (3–5 MΩ) filled with a K^+^-based internal solution containing (in mM): 135 K-gluconate, 5 NaCl, 10 HEPES, 5 MgATP, 0.3 GTP, and 10 phosphocreatine, supplemented with 0.3 % biotin (pH 7.3, 285−290 mOsm). Signals were amplified using a MultiClamp 700B amplifier, digitized at 20 kHz using a Digidata 1550B system (Molecular Devices), and low-pass filtered at 6 kHz during acquisition.

To record evoked inhibitory transmission in voltage clamp mode, the physiological chloride balance was preserved using an internal pipette solution with a low concentration of Cl^-^ ions. Therefore, due to the more negative reversal potential of synaptic GABAergic currents, which was -76±0.60 mV (n = 70), hyperpolarizing inhibitory postsynaptic currents (IPSC) were recorded at a holding potential of -55 mV every 10 s. To determine the reversal potential of synaptic GABA_A_ receptors, IPSCs were recorded at various holding potentials ranging from −90 to −55 mV. Simultaneously, at 0.1 Hz, a 1-second step to −65 mV was applied to record depolarizing excitatory postsynaptic currents (EPSC). Baseline activity was recorded for at least 10 min, followed by measurement of synaptic activity for at least 30 min after the application of NMDA. In these experiments, no pharmacological blockers of synaptic receptors were used. At the end of each recording, pipettes were carefully retracted, re-establishing the GΩ seal and ensuring gentle closure of the cell membrane to preserve its structure and intracellular content. The slices were then fixed and processed for *post hoc* morphological analysis.

All recordings were conducted in postsynaptic interneurons located in the *stratum oriens* of the CA1 hippocampal region. Long-term synaptic plasticity was induced by briefly exposing slices to 20 μM NMDA in the bath solution for 1 or 2 min. The magnitude of GABAergic plasticity was quantified as the percentage change in IPSC amplitude, calculated by comparing the baseline amplitude (averaged over the 6 min preceding NMDA exposure) with the amplitude averaged 26–30 min following NMDA treatment. The series and membrane resistances were evaluated using a -10 mV hyperpolarizing pulse applied every 10 s. Recordings were not corrected for liquid junction potential, and cells were excluded from analysis if the access resistance fluctuated by more than 20 % during the course of the experiment.

To examine the short-term plasticity of IPSCs, pairs of stimuli were delivered at varying inter-stimulus intervals (50 ms, 200 ms, 500 ms, 2 s, and 10 s). The paired-pulse ratio was calculated by dividing the amplitude of the second IPSC by that of the first. To assess burst responses, trains of eight stimuli were applied at 200 ms intervals, and burst depression was quantified as the ratio of the average amplitude of the 6th to 8th responses to the amplitude of the first response within each train.

At the beginning of each experiment, the firing pattern of the postsynaptic interneuron was recorded in current-clamp mode using a series of 0.5-second current steps ranging from -200 pA to 300 pA, incremented at 25 pA intervals. The input resistance was calculated by performing a linear regression on the voltage deflections in response to the applied current steps. The membrane potential at which the slope of the voltage trajectory reached 10 mV/ms was used to define the action potential threshold. The action potential amplitude was measured as the difference between the membrane potential at threshold and the peak of the action potential. Both action potential amplitude and half-width were determined for the first action potential elicited by a depolarizing current pulse (Brzdąk et al., 2023; Wyroślak et al., 2023).

The firing frequency was calculated as the number of action potentials generated during the spike train in response to the strongest depolarizing current pulse. In some interneurons, the injection of hyperpolarizing current pulses elicited a pronounced “sag” response, characteristic of a hyperpolarization-activated cationic current (I_h_), which becomes active during membrane hyperpolarization. To quantify the sag index for each cell, the steady-state voltage deflection was subtracted from the peak negative voltage deflection evoked by the first hyperpolarizing stimulus.

### Optogenetic and electrode stimulation

Stimulation of presynaptic VIP-positive interneurons expressing channelrhodopsin-2 was elicited using 2-ms blue light pulses. Light stimulation was generated using a Lambda DG-4 source (Sutter Instruments) equipped with an HQ470/40x excitation filter (Chroma) and delivered to the sample through a 40×, 1.0 NA objective (Zeiss) centered above the CA1 *stratum oriens*. The aperture of the objective was kept fully open to evoke maximal IPSCs. The power density of the light stimulation was monitored between experiments and remained unchanged throughout and between the recordings.

Excitatory synaptic transmission to CA1 interneurons in the *stratum oriens* was elicited using a tungsten bipolar extracellular stimulating electrode (FHC) positioned in the CA1 *stratum radiatum*, approximately 150 µm from the recorded cell. Presynaptic excitatory axons were excited with 200 µs pulses (50–150 µA) delivered by a constant-current stimulator. In some recordings, stimulation in the *stratum radiatum* induced biphasic EPSCs in postsynaptic CA1 *stratum oriens* interneurons, where the first EPSC was driven by feedforward excitation, followed by a second EPSC resulting from disynaptic feedback excitation. This study specifically focuses on the amplitude of the first EPSC, isolating the feedforward excitatory input to the CA1 interneurons in the *stratum oriens* for further analysis.

### Morphology reconstruction

After electrophysiological recordings, brain slices were fixed overnight in a solution containing 2 % paraformaldehyde and 0.2 % picric acid in phosphate-buffered saline (PBS). The following day, the tissue was rinsed in PBS three times for 10 min each, permeabilized with 0.5 % Triton X-100 in PBS, and blocked using 10 % normal horse serum. Biotin-filled cells were labeled with streptavidin-conjugated Alexa Fluor 488 or Alexa Fluor 568 (#S11223 and #S11226; Thermo Fisher Scientific) for 12 h at 4 °C. The slices were then mounted on Superfrost microscope slides (Thermo Scientific) using Vectashield with DAPI (Sigma/Merck). Confocal images were acquired using an Olympus FluoView 3000 microscope equipped with a 10× objective lens. Cell morphology was reconstructed from Z-stack images obtained at 1-µm intervals along the z-axis.

### Cell identification criteria

We initially attempted to identify the recorded cell types by staining for markers such as parvalbumin (PV), somatostatin (SST), and neuropeptide Y. However, in biotin-labeled neurons from which recordings were obtained, these markers were typically absent, even when morphological identification clearly suggested the cell type (e.g., OLM cells with the expected SST signal). We suspect that this marker washout resulted from the particularly long recordings in the whole-cell configuration (up to 1 hour), which were necessary to characterize long-term plasticity effects. Therefore, we were constrained to rely on strict identification criteria based on morphological and electrophysiological features. We classified all recorded interneurons into two broad categories. Fast-spiking (FS) interneurons were identified based on firing pattern properties, including an action potential (AP) half-width of <0.5 ms, firing frequency >90 Hz, and sag potential <6 mV. The second class of INs in the *stratum oriens* was morphologically classified as O-LM/bistratified cells based on their axonal projections to the *stratum lacunosum-moleculare* or *stratum radiatum*.

Among all recorded neurons with soma located in the *stratum oriens*, five exhibited the distinct morphology of pyramidal cells, including basal and apical dendrites with clearly visible dendritic spines. These cells were excluded from the analysis as they were not interneurons.

### Data analysis and statistics

Electrophysiological recordings were analyzed using the Clampfit 10.7. The onset of IPSCs was characterized by a 10 % – 90 % rise time. Decay kinetics of IPSCs were determined by fitting the biexponential function y(t)=A_1_exp(-t/τ_fast_)+A_2_exp(-t/τ_slow_), where τ_fast_ and τ_slow_ represent time constants, and A_1_ and A_2_ indicate amplitudes of fast and slow components, respectively. The mean decay time constant was calculated as τ_mean_=a_1_τ_fast_+a_2_τ_slow_, where a_1_=A_1_/(A_1_+A_2_), and a_2_=A_2_(A_1_+A_2_).

Data normality was assessed using the Shapiro-Wilk test and variance was evaluated using the Brown-Forsythe test. Paired data were analyzed using a two-tailed paired t-test or Wilcoxon test for non-normally distributed data. Pearson’s rank correlation was used to assess the correlations. Normally distributed data are presented as the mean ± SEM, whereas non-normally distributed data are presented as medians and interquartile ranges. Differences were considered significant when *p* < 0.05, and are indicated in the figures as *p* < 0.001 (∗∗∗), *p* < 0.01 (∗∗), and *p* < 0.05 (∗). Representative traces constitute an average of 30 traces obtained during the 5 min baseline period or during the final 5 min of the recordings. For all LTP experiments, n values represent cells from distinct slices.

## Results

### Diverse properties of VIP-specific inputs onto CA1 *stratum oriens* interneurons

To investigate the properties of I→I synapses formed by VIP-positive presynaptic neurons targeting postsynaptic *stratum oriens* interneurons (*so*INs), we performed single-cell patch-clamp recordings, followed by electrophysiological characterization and morphological reconstruction of the recorded cells. The light-evoked VIP→*so*IN inhibitory postsynaptic currents (IPSC) exhibited an average amplitude of 47.5 ± 3.9 pA (n = 65), a rise time of 3.7 ± 0.16 ms, and a decay time constant (τ_mean_) of 15.8 ± 1.1 ms (Fig. 1A-D).

**Figure 1.**
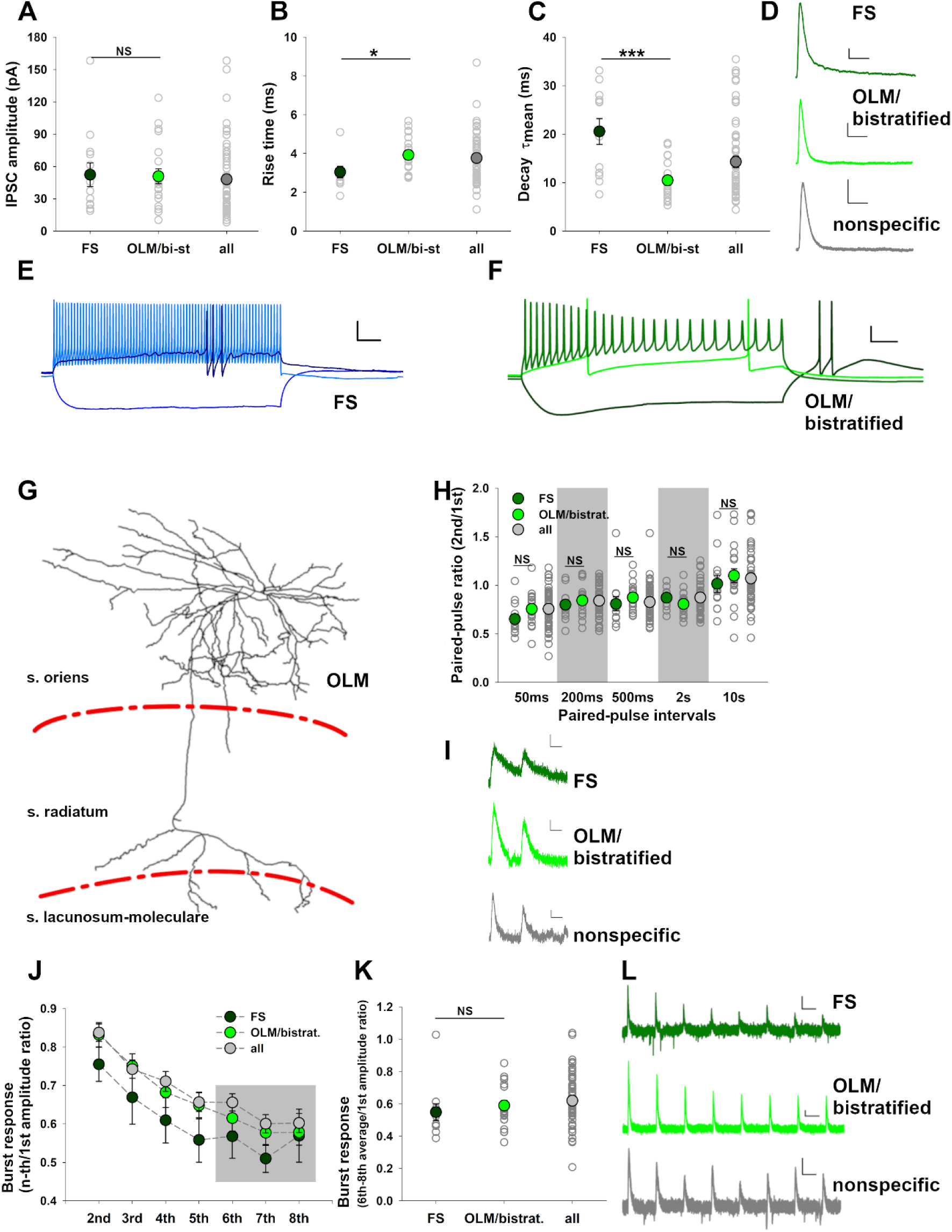
Kinetic properties of VIP-IN-derived IPSCs demonstrate target cell specificity in CA1 stratum oriens interneurons (soINs). **(A)** Summary plot of VIP→soINs IPSC amplitudes recorded from fast-spiking (FS) interneurons (n = 13) and OLM/bistratified interneurons (n = 20). Cumulative data from all recordings, including INs that were not confidently classified into these subgroups, are also shown (n = 65). **(B)** IPSC rise times recorded from FS neurons (n = 11) and OLM/bistratified (n = 20) interneurons, showing shorter rise times in FS cells. Cumulative data from all recorded soINs are shown (n = 49). **(C)** Mean decay time constants (τmean) of VIP→soINs IPSCs recorded from FS (n = 12) and OLM/bistratified (n = 17) interneurons. FS interneurons exhibited significantly prolonged decay times compared to OLM/bistratified cells (cumulative: n = 58). **(D)** Representative traces of IPSCs recorded from FS neurons and OLM/bistratified interneurons in response to light activation of ChR2-expressing VIP interneurons. Scale: 20 pA/30 ms. **(E, F)** Representative firing patterns of FS **(E)** and OLM/bistratified **(F)** interneurons in response to 500-ms current injections (-200 pA, threshold, and +300 pA). FS interneurons displayed high-frequency firing with narrow action potentials, whereas OLM/bistratified interneurons exhibited slower firing and broader action potentials. Scale: 20 mV/50 ms. **(G)** Morphological reconstruction of a biotin-filled OLM interneuron (classified into the OLM/bistratified group), illustrating characteristic axonal projections to the stratum lacunosum-moleculare. **(H)** Paired-pulse ratio (PPR) analysis of VIP-IN-derived IPSCs in FS neurons and OLM/bistratified interneurons, highlighting the lack of cell-type-specific synaptic short-term plasticity. Short-term plasticity was assessed in response to presynaptic paired-pulse stimulation with interpulse intervals of 50 ms, 200 ms, 500 ms, 2 s, and 10 s. **(I)** Exemplary postsynaptic responses recorded in soINs in response to paired-pulse stimulation of VIP interneurons at 50-ms intervals. Scale: 10pA/30ms. **(J)** Burst-induced IPSC depression in all recorded soINs, including FS neurons and OLM/bistratified interneurons. Average ratios of the amplitude of the n^th^ response to the first IPSC within a burst are shown for all analyzed groups. The gray shaded area indicates the responses used to calculate the burst-induced depression index. **(K)** Summary of burst-induced IPSC depression in FS neurons and OLM/bistratified interneurons, showing similar responses between cell types. The burst-induced depression parameter was calculated as the ratio of the mean averaged IPSC amplitude of the 6th–8th responses within a burst (gray area in (E)) to the amplitude of the 1^st^ IPSC. **(L)** Exemplary postsynaptic responses recorded in soINs in response to presynaptic burst stimulation of VIP interneurons with a 200-ms interval. Scale: 10pA/100ms. Data are presented as the mean ± SEM. Statistical significance between the FS and OLM/bistratified groups was assessed using t-tests, with significance levels indicated on each plot (NS – nonsignificant).

Analysis of firing patterns and morphological features allowed for *post hoc* classification of CA1 *stratum oriens* INs into two broad subtypes: fast-spiking (FS) interneurons and OLM/bistratified cells. FS interneurons were characterized by their short half-width of action potentials (AP) (0.37 ± 0.02 ms), minimal or absent sag potential (-2.2 ± 0.5 mV at -200 pA holding current) and high maximum firing frequency (103 ± 11.5 Hz; Fig. 1E, Table 1). In contrast, OLM/bistratified interneurons were identified primarily based on their axonal morphology, which extended to the *stratum lacunosum-moleculare* or *radiatum* (Fig. 1F,G). Their electrophysiological properties differed significantly from FS interneurons, including broader AP half-widths (0.61 ± 0.05 ms; *p* < 0.001 vs. the FS group), pronounced sag potentials (-11.6 ± 1.3 mV at -200 pA holding current; *p* < 0.001), and lower maximum firing frequencies (50.5 ± 4.1 Hz; *p* < 0.001; Table 1). Although the electrophysiological properties of CA1 interneurons form a continuous distribution, the AP half-width has been reported to be a reliable distinguishing feature between PV-positive fast-spiking basket cells and other interneuron types (Tricoire et al., 2011).

**Table 1.** Electrophysiological properties of interneuron groups identified in the CA1 stratum oriens. Significance was assessed using a paired t-test.

| Parameter | FS (n = 12) | OLM/Bistratified (n = 20) | p-value |
| --- | --- | --- | --- |
| Resting membrane potential (mV) | -64.3 ± 2.1 | -64.3 ± 0.97 | 0.996 |
| Maximum firing frequency (Hz) | 103.3 ± 11.5 | 50.5 ± 4.1 | 1.59 × 10<sup>-5</sup> |
| Sag potential (mV) | -2.2 ± 1.4 | -11.6 ± 1.3 | 3.7 × 10<sup>-6</sup> |
| AP half-width (ms) | 0.37 ± 0.02 | 0.62 ± 0.05 | 0.0003 |
| Firing threshold (mV) | -33.1 ± 1.5 | -38.8 ± 1.2 | 0.005 |
| Membrane resistance (MΩ) | 119.7 ± 10 | 303.3 ± 35.1 | 0.0003 |
| AP amplitude (mV) | 62.3 ± 3.1 | 76.4 ± 1.9 | 0.0003 |

Optogenetically evoked IPSCs mediated by VIP input revealed distinct kinetic differences between the two subtypes of postsynaptic interneurons. In FS cells, IPSCs had shorter rise times (3.05 ± 0.3 ms Fig. 1B) and longer decay times (τ_mean_: 20.6 ± 2.7 ms; Fig. 1C, D). Interestingly, the τ_mean_ in FS cells showed substantial variability, which did not correlate with any other measured parameter (data not shown). This suggests that the variability is likely intrinsic at the synaptic level rather than reflecting differences between cellular subtypes within the FS group. In contrast, VIP-derived IPSCs in OLM/bistratified INs exhibited longer rise times (3.9 ± 0.2 ms; *p* = 0.022 vs. FS cells, Fig. 1B) but shorter decay times (τ_mean_: 10.5 ± 1.0 ms; *p* = 0.0005 vs. FS cells, Fig. 1C, D). The decay time constants observed in OLM cells are consistent with previous findings from paired recordings by Tyan et al., (2014). These differences in IPSC kinetics suggest that VIP interneurons form GABAergic synapses with distinct properties on these two classes of *so*INs, and our next step was to investigate whether these differences are reflected in short-and long-term plasticity.

### Short-term plasticity in VIP→*so*IN synapses

Short-term plasticity in inhibitory synapses (iSTP) plays a critical role in synaptic function and circuit dynamics (Anwar et al., 2017). Previous studies have demonstrated input-specific iSTP of IPSCs in CA1 pyramidal neurons (Udakis et al., 2020; Wiera et al., 2024). Thus, examining this type of plasticity in the projections from VIP-positive interneurons to *so*INs is of particular interest. Paired-pulse stimulation revealed striking variability in iSTP, ranging from profound short-term depression (iSTD, with up to an 80 % decrease in IPSC amplitude), to no detectable plasticity, and even short-term potentiation, which was most prominent at longer intervals between pulses (e.g., 10 s). For all recorded *so*INs combined, including cases without successful reconstruction or those not classified as FS or OLM/bistratified cells, paired-pulse ratios showed an overall trend of iSTD across most tested intervals, with facilitation observed only at the 10 s interval (paired-pulse ratios; 50 ms: 0.76 ± 0.025; 200 ms: 0.84 ± 0.02; 500 ms: 0.83 ± 0.022; 2 s: 0.87 ± 0.02; 10 s: 1.07 ± 0.04; Fig. 1H). Comparisons between FS-INs and OLM/bistratified interneurons revealed no statistically significant differences in the response to paired-pulse stimulation. FS interneurons exhibited a tendency towards stronger iSTD at shorter intervals, although these differences were not statistically significant (e.g., 50 ms: FS 0.65 ± 0.05 vs. OLM/bistratified 0.75 ± 0.04, *p* = 0.11; 500 ms: FS 0.8 ± 0.07 vs. OLM/bistratified 0.87 ± 0.03, *p* = 0.36; Fig. 1H).

In response to burst stimulation, cumulative data from all recorded cells showed an average 40 % reduction in synaptic response amplitude (0.62 ± 0.022; Fig. 1I-K). No significant differences were observed between FS-INs and OLM/bistratified cells in burst-induced depression (FS: 0.55 ± 0.052; OLM/bistratified: 0.59 ± 0.031; *p* = 0.48; Fig. 1I, J). Thus, despite clear differences in IPSC kinetics between FS and OLM/bistratified cells (Fig. 1A-D), their short-term plasticity profiles were remarkably similar. These findings suggest that while VIP inputs to FS neurons and OLM/bistratified cells differ in synaptic kinetics, they do not exhibit subtype-specific short-term plasticity. Instead, the observed variability in iSTP likely reflects the functional heterogeneity of VIP→*so*IN synapses rather than differences between the targeted interneuron subtypes.

### VIP→*so*IN synapses show NMDA-dependent GABAergic plasticity

Previous studies have shown that long-term excitatory plasticity in pyramidal neurons is frequently accompanied by long-term plastic changes at inhibitory synapses on the same cell (Chiu et al., 2018; Field et al., 2020; Wiera et al., 2024). To determine whether such co-expression of excitatory and inhibitory plasticity occurs in *so*INs, we used an NMDA application protocol previously shown to induce plasticity in both excitatory and inhibitory synapses in pyramidal neurons (Wiera et al., 2024) and interneurons (Brzdąk et al., 2023). Because NMDA-induced plasticity is highly sensitive to the timing of receptor activation (Wiera et al. 2024), we tested two application durations. We began with a 1-minute NMDA application to assess its effects on IPSCs recorded from *so*INs receiving input from VIP-positive INs. In OLM/bistratified interneurons, this treatment resulted in a significant decrease in IPSC amplitude (before NMDA: 41.4 ± 7.6 pA; 26-30 minutes after: 33.5 ± 5.6 pA; *p* = 0.01; Fig. 2A-C). However, no such changes were observed in fast-spiking (FS) interneurons (before: 52.4 ± 9.7 pA; after: 49.9 ± 12.0 pA; *p* = 0.52; Fig. 2C). When all recorded *so*INs, including unidentified interneurons, were pooled for analysis, the 1-minute application of NMDA led to a long-term reduction in IPSC amplitude, with an average decrease to 85.3 ± 2.7 % of baseline (before: 42 ± 4.0 pA; after: 37.0 ± 4.0 pA; *p* = 0.0002; Fig. 2A-C).

**Figure 2.**
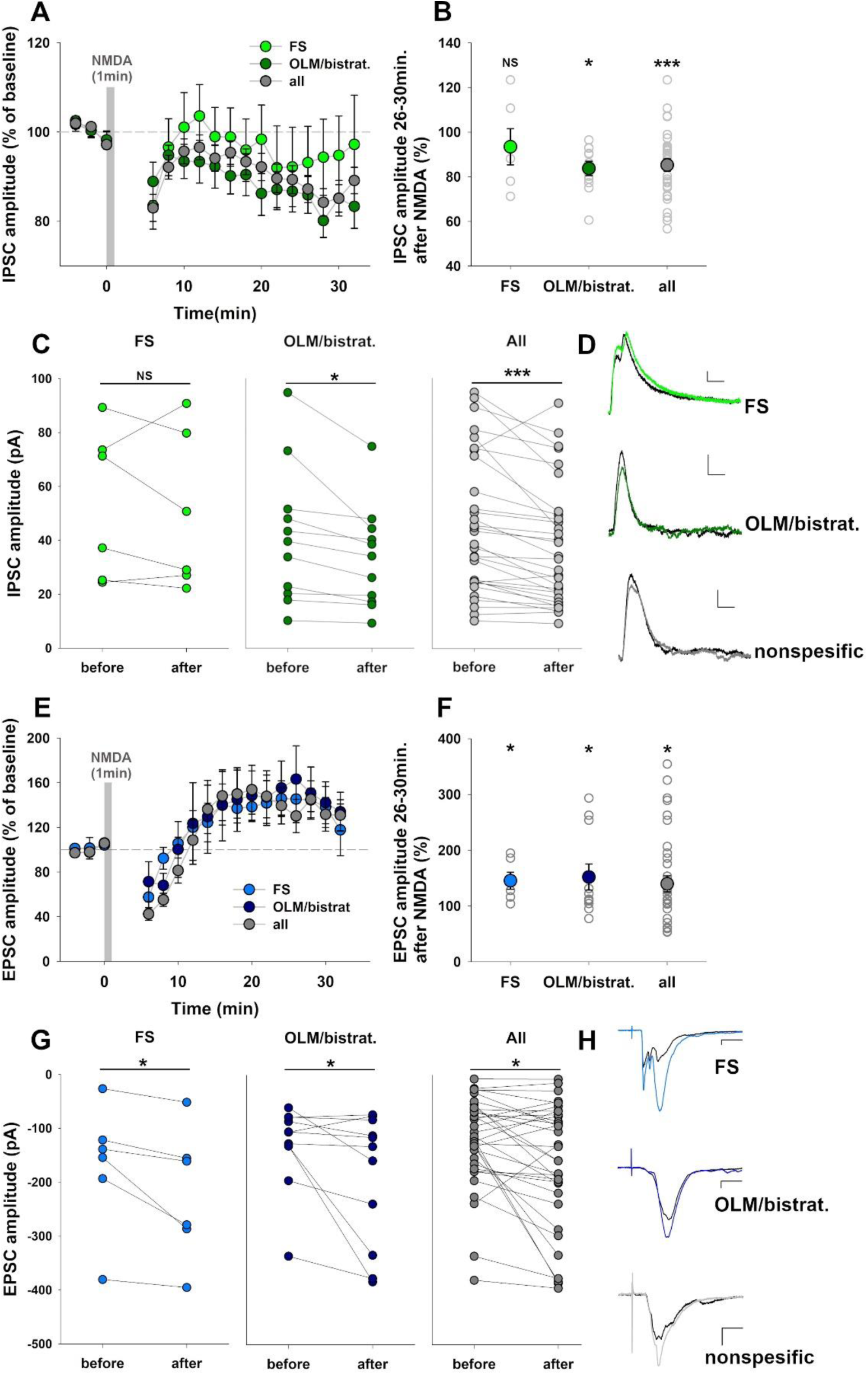
Postsynaptic neuron-specific excitatory-inhibitory coplasticity induced by NMDA application for 1 min in CA1 stratum oriens interneurons (soINs). **(A-D) Long-term inhibitory plasticity at VIP→soIN** synapses **following a 1-minute NMDA treatment**. **(A)** Time course of IPSC amplitude changes after NMDA application in CA1 stratum oriens interneurons. **(B)** Normalized percentage changes in IPSC amplitudes across the experimental groups. **(C)** Summary plot of IPSC amplitude changes (inhibitory LTD or lack of average plasticity) measured before and 26–30 minutes after NMDA treatment. **(D)** Representative IPSC traces recorded before (black) and 30 min after (colored) NMDA treatment, scale 10 pA/10ms. Inhibitory synaptic transmission was evoked by local optogenetic stimulation of VIP+ INs. (**E-H**) Long-term plasticity in excitatory synapses onto CA1 stratum oriens INs induced in response to 1-minute NMDA application, recorded simultaneously with the results presented in (A). Excitatory synaptic transmission was evoked by placing a stimulation electrode in the CA1 stratum radiatum. **(E)** Time course of excitatory postsynaptic current (EPSC) amplitude changes following 1-minute NMDA application in CA1 stratum oriens interneurons. **(F)** Summary plot of EPSC amplitude changes (excitatory LTP) measured 26–30 minutes after NMDA treatment. **(G)** EPSC amplitudes recorded before and 26-30 minutes after NMDA treatment in each group. **(H)** Representative EPSC traces recorded before (black) and 30 min after (colored) NMDA treatment, scale 20 pA/10ms. Sample sizes: FS interneurons (n = 6), OLM/bistratified interneurons (n = 11), and all soINs (including both previous groups; n = 34). Statistical significance between the FS and OLM/bistratified groups was assessed using unpaired t-tests (B,F), while the effect of NMDA treatment was evaluated using paired t-tests (C,G). Significance levels are indicated on each plot.

Paired-pulse stimulation, analyzed before and 30 min after NMDA treatment, provides valuable insights into the locus of plastic changes. Across all recorded *so*INs, the paired-pulse ratio (PPR) at the 50 ms interval increased significantly after the application of NMDA, while no changes were observed at longer stimulation intervals (Fig. S1A). A similar trend was observed in OLM/bistratified INs, which showed a less pronounced paired-pulse depression at the 50 ms interval after NMDA-induced iLTD (Fig. S1B). In contrast, FS neurons did not show changes in PPR after NMDA treatment (Fig. S1C). These results indicate that NMDA stimulation had a modest impact on iSTP in OLM/bistratified INs by increasing the PPR, but this effect was limited to a brief time window (50 ms).

To identify potential mechanisms influencing the induction and expression of VIP→*so*IN iLTD, we investigated correlations between the extent of iLTD and various electrophysiological properties, IPSC kinetics, and short-term inhibitory plasticity. Interestingly, no significant correlations were found between iLTD magnitude and either the age of the mice (Fig. S2A) or the active membrane properties of postsynaptic *so*INs (Fig. S2B-G). However, a significant positive correlation emerged between the initial reversal potential of VIP→*so*IN IPSCs and the magnitude of inhibitory plasticity. Specifically, more depolarized GABA reversal potentials were associated with a reduced extent of iLTD (Pearson coefficient = 0.56, *p* = 0.0013; Fig. S2F). In contrast, we found no significant correlations between the magnitude of the iLTD and the rise time, decay time, or paired-pulse ratios of the IPSC (Fig. S2H-N). Additionally, stronger burst-induced depression correlated with more pronounced iLTD at VIP→*so*IN synapses following NMDA application (Pearson’s coefficient = 0.38, *p* = 0.04; Fig. S2O). Together, these findings indicate that the mechanisms regulating the extent of inhibitory plasticity at VIP→*so*IN triggered by a 1-minute NMDA application may depend on the electrical driving force at GABAergic synapses (reversal potential) and are partially correlated with initial short-term plasticity.

Taking advantage of our experimental design, we simultaneously recorded IPSCs and EPSCs to explore their parallel plastic changes. Following a 1-minute NMDA treatment, we observed a consistent and significant increase in EPSC amplitude, indicative of excitatory long-term potentiation (eLTP). Across all recorded *so*INs, EPSC amplitude increased to 146.8 ± 13.2 % of baseline (before: -121.5 ± 13.7 pA; after: -188.4 ± 18.9 pA; *p* = 0.002; Fig. 2E-G). This eLTP was evident in both interneuron subtypes. FS interneurons showed a 145.5 ± 15.2 % increase (before: -148.3 ± 45.61 pA; after: -221.6 ± 49.9 pA; *p* = 0.04), while OLM/bistratified interneurons exhibited a 152 ± 23.4 % increase (before: -129.5 ± 23.5 pA; after: -189.7 ± 36.9 pA; *p* = 0.05; Fig. 2E-G). These results reveal a striking divergence in the co-expression of plasticity between the two IN subtypes. OLM/bistratified INs displayed asymmetric coplasticity, characterized by eLTP at their excitatory inputs and iLTD at their inhibitory VIP-positive inputs. In contrast, FS interneurons demonstrated eLTP without significant changes in their VIP-mediated inhibitory drive. This subtype-specific dissociation underscores the dependence of coplasticity patterns on the interneuron subtype and their functional roles within hippocampal circuits.

### Co-expression of excitatory and inhibitory plasticity in VIP→*so*IN synapses following 2-minute NMDA application

To explore how prolonged activation of NMDA receptors affects the balance between excitatory and inhibitory plasticity, we extended the application of NMDA to 2 min and assessed its effects on VIP-mediated IPSCs and EPSCs in the *so*INs (Fig. 3 A-B). This longer NMDA exposure led to a consistent reduction in IPSC amplitudes across all recorded *so*INs, indicative of iLTD. IPSC amplitudes decreased to 83.3 ± 3.2 % of baseline for all recorded cells (before: 51.0 ± 6.9 pA; after: 43.1 ± 6.5 pA; *p* = 0.0004; Fig. 3A-C). This reduction was evident across both interneuron subtypes. Fast-spiking (FS) interneurons showed a significant decrease in IPSC amplitude (81.4 ± 4.5 %; before: 56.7 ± 26.6 pA; after: 47.6 ± 23.7 pA; *p* = 0.045), as did OLM/bistratified interneurons (81.0 ± 3.4 %; before: 58.6 ± 12.5 pA; after: 48.6 ± 11.2 pA; *p* = 0.006; Fig. 3A-C). Analysis of paired-pulse ratios for IPSCs (for all *so*INs) following 2-minute NMDA application revealed no significant changes (S1D-F), suggesting that the observed iLTD was mediated postsynaptically. Furthermore, no significant correlations were identified between the magnitude of VIP→*so*IN iLTD and factors such as active membrane properties of postsynaptic neurons, IPSC kinetics, short-term plasticity, animal age, or GABAergic reversal potential (Fig. S3).

**Figure 3.**
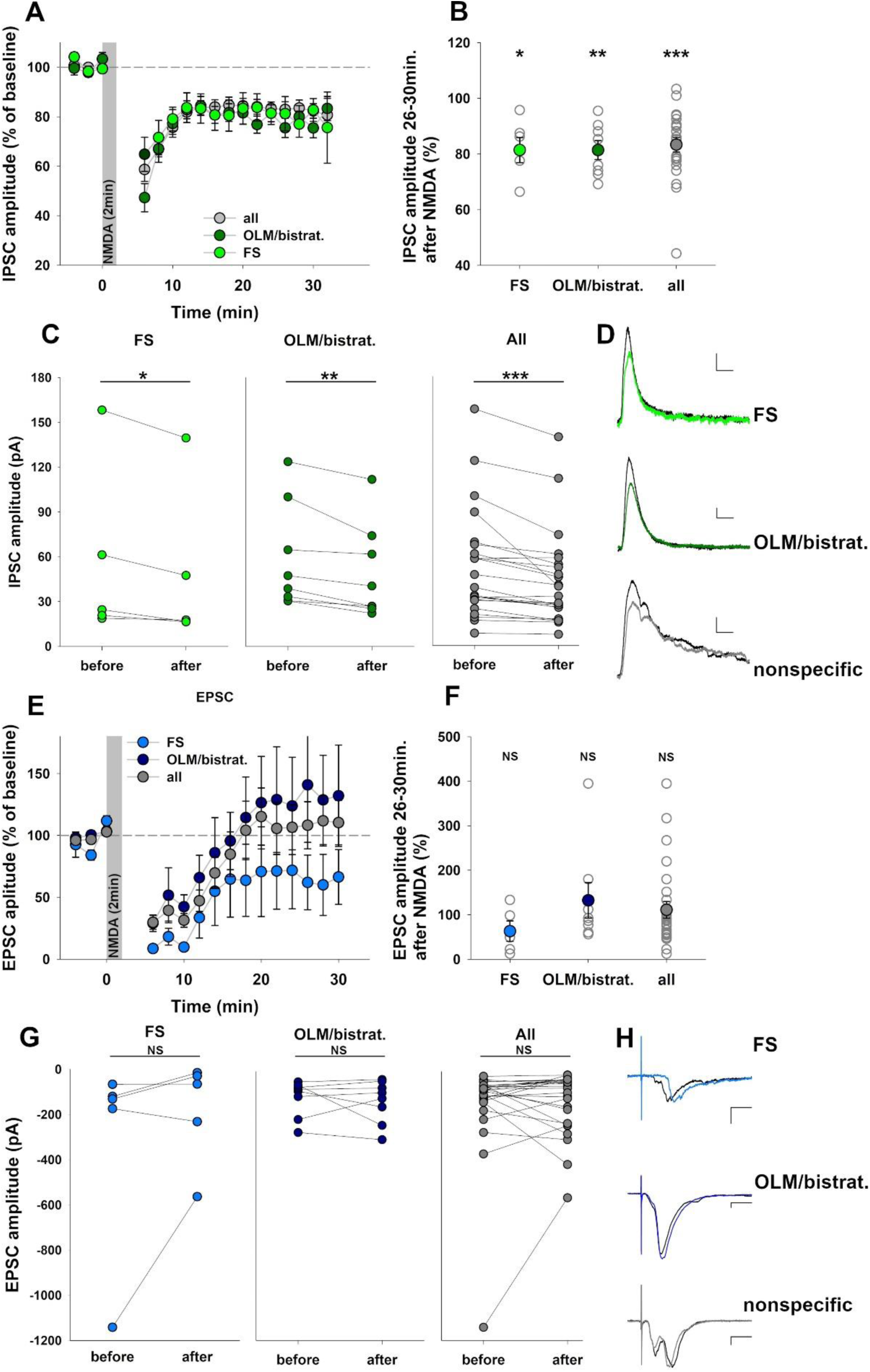
Prolonged 2-minute NMDA application induces iLTD of VIP-derived IPSCs in stratum oriens interneurons. **(A-D)** Long-term inhibitory plasticity at VIP→soIN synapses in response to a 2-minute NMDA treatment. **(A)** Time course of IPSC amplitude changes in CA1 stratum oriens interneurons induced by NMDA application (experimental groups: all recorded INs, fast-spiking INs (FS) and OLM/bistratified INs). **(B)** Normalized percentage changes in IPSC amplitudes (iLTD) across experimental groups. **(C)** Comparison of IPSC amplitudes before and 26-30 minutes after NMDA treatment in individual recorded cells within each group. **(D)** Representative IPSC traces recorded before (black) and 30 min after (colored) NMDA treatment, scale 10pA/10ms. Inhibitory synaptic transmission was evoked by local optogenetic stimulation of VIP + INs. **(E-H)** Long-term plasticity in excitatory synapses onto stratum oriens INs, induced simultaneously with inhibitory plasticity by a 2-minute NMDA application. Excitatory synaptic transmission was evoked by placing a stimulation electrode in the CA1 stratum radiatum. **(E)** Time course of EPSC amplitude changes following NMDA treatment. **(F)** Normalized percentage changes in EPSC amplitudes across the experimental groups. **(G)** Comparison of EPSC amplitudes recorded before and 26-30 minutes after NMDA treatment in each group. **(H)** Representative EPSC traces recorded before (black) and 30 min after (colored) NMDA application, scale 20pA/10ms. Sample sizes: FS interneurons (n = 5), OLM/bistratified interneurons (n = 8), all recorded soINs combined (n = 24). Statistical significance between the FS and OLM/bistratified groups was assessed using unpaired t-tests (B, F), whereas the effect of NMDA treatment was evaluated using paired t-tests (C, G). Significance levels are indicated on each plot.

Interestingly, no excitatory LTP was observed for the longer NMDA application protocol. Across all recordings, EPSC amplitudes showed no significant change following the 2-minute NMDA application (111.1 ± 18.7 %; before: 144.4 ± 42.8 pA; after: 133.2 ± 27.3 pA; *p* = 0.7; Fig. 3E-G). This lack of significant plasticity was observed for OLM/bistratified interneurons, which showed no changes in EPSC amplitudes (132.4 ± 39.9 %; before: -117.5 ± 28.7 pA; after: -133.6 ± 33.8 pA; *p* = 0.59; Fig. 3F-G). FS interneurons showed a slight trend towards excitatory eLTD, with EPSC amplitudes decreasing to 63.2 ± 23.0 % of baseline (before: 326.8 ± 204.4 pA; after: 181.1 ± 103.1 pA; *p* = 0.26; Fig. 3F-G), but this effect did not reach statistical significance, likely due to the high variability in EPSC measurements. Notably, comparisons of the magnitude of excitatory plasticity between FS and OLM/bistratified interneurons also revealed no significant differences (*p* = 0.23; Fig. 3F). These findings indicate that extending the duration of NMDA application alters the balance of co-expressed plasticity in *so*INs compared with shorter NMDA exposure. Although shorter NMDA treatments induce both excitatory and inhibitory plasticity, prolonged exposure seems to predominantly drive inhibitory plasticity, with excitatory inputs showing minimal or no change.

### Relationship between inhibitory and excitatory plasticity in *stratum oriens* interneurons

To investigate the co-expression of inhibitory and excitatory plasticity within individual *stratum oriens* interneurons, we analyzed the correlation between percentage changes in IPSC and EPSC amplitudes in response to NMDA application across all recorded cells. Scatter plots revealed a clear change in coplasticity patterns as the duration of exposure to NMDA increased. Considering all recorded *so*INs, after a 1-minute NMDA application, *so*INs exhibited opposing and asymmetric plasticity, with I→I iLTD at VIP inputs and eLTP at E→I excitatory inputs (Fig. 4A). In contrast, a 2-minute NMDA treatment induced a distinct shift, while excitatory inputs showed only minor and statistically insignificant changes, inhibitory inputs showed highly significant I→I iLTD at VIP-mediated synapses (Fig. 4A). This pattern was consistently observed in identified postsynaptic OLM/bistratified interneurons (Fig. 4C). The trends in FS cells were less defined. However, a direct comparison of plasticity profiles between the two NMDA exposure durations revealed significant shifts. Prolonged NMDA application led to the emergence of robust iLTD at GABAergic I→I synapses, transitioning from a lack of inhibitory plasticity for 1 min NMDA pulses. Simultaneously, excitatory plasticity shifted from LTP towards a marked reduction in EPSC amplitude changes (Fig. 4B, Fig. S4).

**Figure 4.**
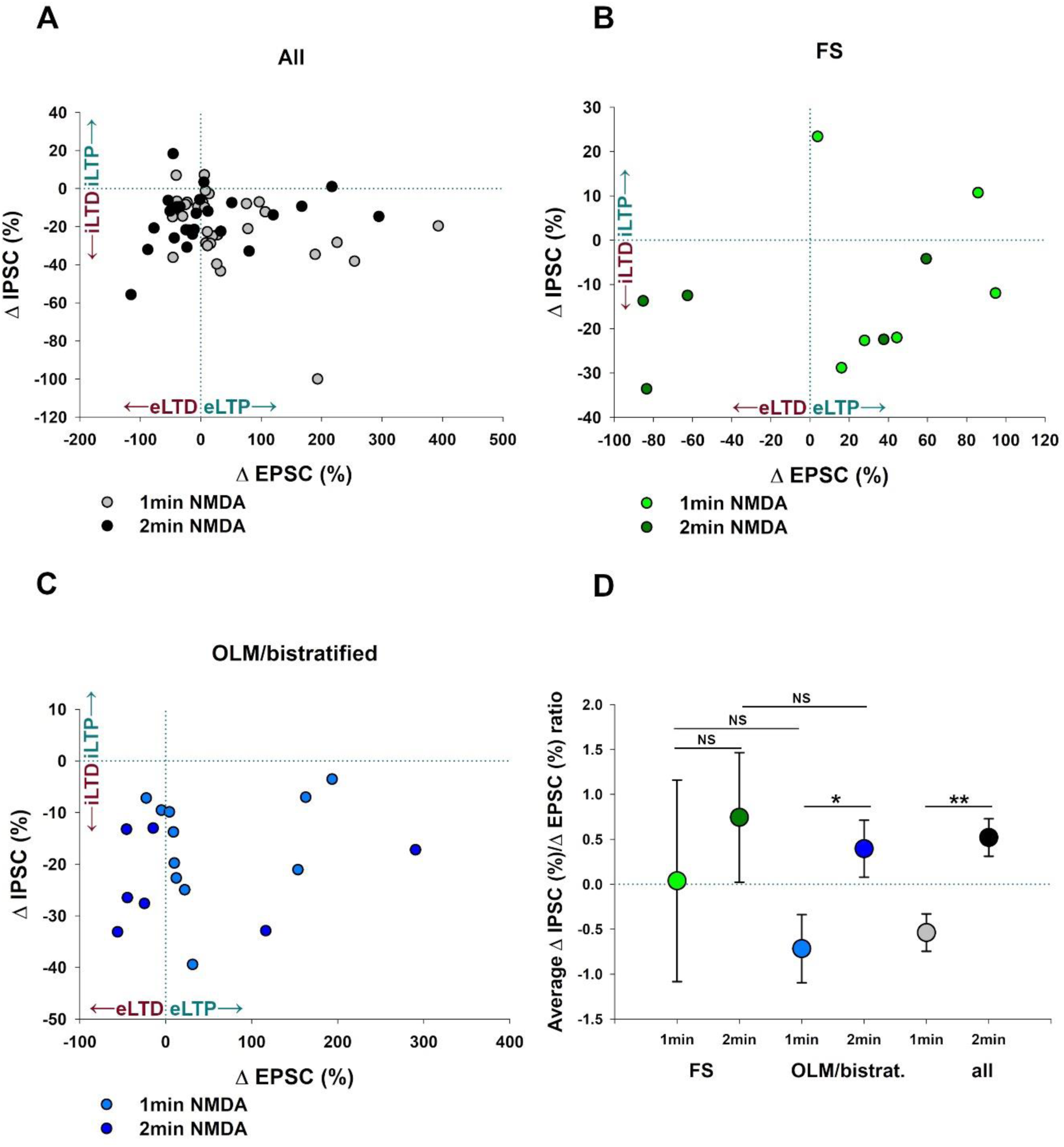
Altering NMDA exposure duration switches the balance between inhibitory and excitatory plasticity co-expressed in CA1 stratum oriens interneurons. **(A-C)** Scatter plots illustrating the direction and magnitude of excitatory and VIP-derived inhibitory long-term plasticity in individual CA1 stratum oriens interneurons (soINs) induced by 1-minute or 2-minute NMDA application. The ΔEPSC and ΔIPSC parameters show changes in the amplitudes of EPSC and IPSC, respectively, after NMDA-induced plasticity. Positive values indicate LTP, negative ones -LTD **(A)** Cumulative data showing plasticity across all individual recorded soINs following 1-minute NMDA treatment (n = 30) or 2-minute NMDA treatment (n = 25). **(B)** Coplasticity expressed in fast-spiking interneurons (FS) recorded after 1-minute (n = 6) or 2-minute NMDA treatment (n = 5). **(C)** Coplasticity expressed in OLM/bistratified interneurons induced by 1-minute (n = 11) or 2-minute NMDA treatment (n = 7). **(D)** Statistics of the ratio of inhibitory plasticity (VIP→soINs) to the magnitude of simultaneously co-expressed excitatory plasticity induced by NMDA application for 1 or 2 minutes in postsynaptic CA1 stratum oriens interneurons. The inhibitory-to-excitatory plasticity ratio was calculated by dividing ΔIPSC/ΔEPSC (see Results section). Statistical significance was assessed using unpaired t-tests, with the significance levels indicated on each plot.

To quantify the effects of NMDA exposure on inhibitory/excitatory balance, we calculated the ratio of percentage changes in IPSC and EPSC amplitudes for each cell. This ratio, defined as ΔIPSC/ΔEPSC, reflects the relationship between concurrent plasticity in inhibitory and excitatory synapses. Positive ΔIPSC and ΔEPSC values indicate synaptic potentiation (iLTP and eLTP, respectively), whereas negative values represent long-term depression (iLTD and eLTD). A positive ratio indicates symmetrical plasticity, where both inputs undergo similar changes (e.g., co-expression of iLTP and eLTP or iLTD and eLTD), whereas a negative ratio indicates asymmetrical plasticity, where LTP in one pathway coincides with LTD or lack of plasticity in the other.

After 1 minute of NMDA application, the average ΔIPSC/ΔEPSC ratio for all recorded *so*INs was negative (-0.53 ± 0.21; Fig. 4D), reflecting a predominance of cells in the lower right quadrant of the scatter plot (Fig. 4A). Extending NMDA exposure for 2 min shifted this balance towards symmetrical plasticity, with the majority of INs exhibiting long-term depression in both inputs, reflecting a clustering of cells in the lower-left quadrant of the scatter plot (Figs. 4A and D). For 2-minute NMDA application, the ΔIPSC/ΔEPSC ratio became positive (0.52 ± 0.21), and was significantly different compared to 1-minute (*p* = 0.001; Fig. 4D). This trend was particularly evident in OLM/bistratified interneurons, where statistical significance in ΔIPSC/ΔEPSC ratio between NMDA application durations was observed (1-minute NMDA: -0.72 ± 0.38, 2-minute NMDA: 0.39 ± 0.31; *p* = 0.043; Fig. 4C, D). In contrast, for fast-spiking (FS) interneurons, the extended NMDA exposure did not result in a significant change in the ΔIPSC/ΔEPSC ratio (1-minute NMDA: 0.034 ± 1.12, 2-minute NMDA: 0.74 ± 0.73; *p* = 0.20, Mann-Whitney test; Fig. 4D). Nevertheless, prolonged NMDA application revealed a shift in the coplasticity profile, transitioning from the absence of GABAergic plasticity paired with E→I eLTP to pronounced iLTD accompanied by a lack of significant excitatory plasticity. This shift highlights the pivotal role of NMDA receptor activation timing in modulating the plasticity of VIP→*so*IN synapses, producing distinct effects across various postsynaptic interneuron subtypes.

## Discussion

This study examined VIP IN-mediated inhibitory inputs to *stratum oriens* interneurons (VIP→*so*IN) in the CA1 region, focusing on synaptic transmission, I→I plasticity, and NMDA-dependent coplasticity. The major finding of this study is that brief application of NMDA induced plasticity at both inhibitory (VIP input) and excitatory synapses at the INs of interest in a cell-specific manner. Moreover, prolongation of NMDA administration (from 1 to 2 min) altered the relationship between inhibitory and excitatory plasticity in FS and OLM/bistratified INs, indicating a potential impact of these changes on the E/I balance.

### Divergence in synaptic properties and short-term plasticity in distinct VIP→*so*IN synapses

We observed that the two primary interneuron subtypes, fast-spiking (FS) interneurons and OLM/bistratified cells, exhibit distinct IPSC kinetic properties in response to VIP input stimulation. Fast-spiking neurons, which are known to include basket and chandelier cells, exhibited relatively long IPSC decay times and fast rise times, whereas OLM/bistratified interneurons exhibited shorter IPSC decay times and longer rise times (Fig. 1). Notably, the difference in rise times, which may be influenced by dendritic electrotonic properties (Goswami et al., 2012; Wiera et al., 2024), suggests that VIP inhibitory synapses on fast-spiking interneurons are closer to the cell body than those formed on OLM/bistratified cells. In a recent study, McFarlan et al. (2024a) documented similar cell-specific properties of VIP-positive I→I inputs in the motor cortex, showing that VIP-driven IPSCs differ notably between Martinotti and basket cells, particularly in terms of rise time and latency. Specifically, VIP input to cortical parvalbumin-positive basket cells exhibited faster onset kinetics than the same input onto somatostatin-positive Martinotti interneurons (McFarlan et al., 2024a). It is thus noteworthy that although McFarlan et al. (2024a) studied a different model (motor cortex), their observations concerning IPSC onset kinetics are similar to those seen in our recordings from hippocampal CA1 *stratum oriens* interneurons (FS INs comprise mainly PV basket cells, and OLM/bistratified INs are SST-positive as Martinotti cells).

It is particularly intriguing that IPSC decay kinetics in VIP input to fast-spiking interneurons was markedly longer compared to that in OLM/bistratified cells. This suggests that presynaptic VIP interneurons form inhibitory synapses with distinct properties and/or GABA_A_R subunit compositions, a mechanism that has already been described for different inhibitory inputs to the *stratum oriens* somatostatin-positive interneurons (Magnin et al., 2019). Three distinct classes of interneuron-selective interneurons (ISI) have been identified in the hippocampus, two of which, ISI-2 and ISI-3, express VIP. The ISI-2 class predominantly establishes synaptic connections with specific interneurons while avoiding pyramidal cells and interneurons expressing parvalbumin as postsynaptic partners (Pelkey et al., 2017). In contrast, the ISI-3 class primarily targets OLM interneurons but also forms connections with other interneurons, including bistratified cells and PV-positive basket cells, although less frequently (Chamberland and Topolnik, 2012; Tyan et al., 2014). Therefore, the differences in IPSC decay kinetics between VIP→FS and VIP→OLM/bistratified synapses may be due to two distinct classes of VIP interneurons, each targeting different subtypes of interneurons and forming I→I synapses with distinct properties. In our recent study (Brzdąk et al. 2023), we found that mIPSC decay in PV-positive INs was slower than in SST INs, confirming the trend described in the present work for IPSCs in FS and OLM/bistratified neurons. However, it should be considered that, for mIPSCs, synaptic inputs remain unspecified and that the experimental conditions used in the present study were different from those applied by Brzdąk et al. 2023 (see also below).

Interestingly, while we observed significant differences in IPSC kinetics for VIP inputs (e.g., rise and decay times) between FS neurons and OLM/bistratified cells, we did not find clear differences in IPSC amplitude or short-term plasticity across these postsynaptic cell types. This may be due to the heterogeneity of both the VIP input and postsynaptic target populations, which complicates the interpretation of paired-pulse ratio data. Since short-term plasticity is primarily governed by presynaptic properties, our results suggest that the VIP inputs stimulated in our experiments exhibit a similar profile of short-term depression, regardless of the postsynaptic interneuron type. A similar pattern was observed for VIP synapses in the motor cortex (McFarlan et al., 2024a).

### NMDA-induced inhibitory plasticity

Our results demonstrate target cell-specific GABAergic long-term plasticity at VIP→*so*IN synapses in response to NMDA receptor activation. Brief NMDA treatment (1 min) reliably induced inhibitory long-term depression (iLTD) at VIP→OLM/bistratified synapses, but no significant plasticity was observed at VIP→FS connections (Fig. 2). Interestingly, when NMDA application was prolonged to 2 min, robust iLTD was induced in both types of synapses. These findings reveal a fundamental difference between the plasticity at VIP→*so*IN synapses (I→I) compared to that at inhibitory synapses targeting pyramidal cells (I→E synapses). Indeed, our previous studies have shown that a 1-minute NMDA application induced inhibitory iLTP at both SST→PC and PV→PC synapses, whereas longer NMDA exposure continued to elicit iLTP in SST→PC connections but failed to induce any consistent plasticity at PV→PC synapses (Jabłońska et al., 2024; Wiera et al., 2024). Thus, our findings suggest a potential network mechanism by which OLM/bistratified interneurons undergo simultaneous plasticity at their inhibitory input and output synapses. Specifically, iLTD at their VIP-mediated inhibitory inputs is coupled with iLTP at their inhibitory synapses located onto CA1 pyramidal cells (Wiera et al., 2024). This dual plasticity would produce a two-step enhancement of the inhibitory drive onto excitatory cells. At the same time the inhibitory drive to interneurons would weaken because of iLTD occurring at VIP→*so*INs. This diverging inhibitory plasticity at inhibitory and excitatory target neurons is expected to create a dynamic shift in the excitation-inhibition balance that promotes overall network stability (Sadeh and Clopath, 2021).

Observed here differences in the inhibitory plasticity for VIP INs projections onto FS or OLM/bistratified INs are not surprising. In the afore mentioned work of McFarlan et al. (2024b) they demonstrated in the motor cortex that causal pairing induced iLTD in VIP synapses onto Martinotti cells, but not in VIP input to basket cells, highlighting the specificity of plasticity mechanisms at different VIP→IN synapses.

In a recent study Brzdąk et al. (2023) studied NMDA-induced GABAergic plasticity of mIPSCs in SST-and PV-positive CA1 hippocampal interneurons and reported iLTP in the former and iLTD in the latter INs. At first glance, these data appear to contradict our present results for OLM/bistratified interneurons (which are SST-positive) for which iLTD was reported. However, this apparent discrepancy may be explained by differences in the experimental approaches. In the current study, excitatory synaptic transmission was not pharmacologically blocked, the pipette internal solution contained a physiological chloride concentration that produced hyperpolarizing inhibitory currents, and we analyzed IPSC evoked by a specific inhibitory input rather than mIPSCs, which originate from a mix of GABAergic drives. These methodological differences can significantly influence the observed direction of plasticity. Thus, our findings and those of Brzdąk et al. (2023) suggest that different inhibitory inputs to SST-positive interneurons may exhibit distinct direction and mechanisms of NMDA-dependent plasticity.

### Network implications of synaptic coplasticity in CA1 *stratum oriens* interneurons

An important aspect of our study was the simultaneous examination of both inhibitory and excitatory plasticity co-expressed in postsynaptic *so*INs in response to activation of NMDA receptors. Our findings reveal distinct patterns of synaptic coplasticity between OLM/bistratified interneurons and fast-spiking neurons and their different dependence on NMDA transient duration (Figs. 2-4). In OLM/bistratified cells, NMDA application led to opposing asymmetric coplasticity: eLTP (or no plasticity) at excitatory synapses and iLTD at VIP-mediated inhibitory input. This asymmetric modulation of the E/I balance in favor of excitation is likely to enhance the ability of OLM/bistratified cells to regulate hippocampal network activity, particularly by stronger engagement in feedback controlling the dendritic associativity of pyramidal cells (Müller and Remy, 2014; Udakis et al., 2020). In contrast, coplasticity in fast-spiking neurons showed a clear dependence on the duration of NMDA receptor activation. A 1-minute application of NMDA resulted in asymmetric plasticity, with eLTP at excitatory inputs and no changes in GABAergic inputs. However, a 2-minute NMDA application induced iLTD in VIP-mediated inputs while leaving excitatory inputs unchanged. These results underscore the importance of the duration of NMDA receptor activation in modulating the balance between excitation and inhibition, highlighting thus heterosynaptic nature of plastic changes in these circuits. Moreover, in FS neurons, which are primarily responsible for rapid feedforward inhibition (Hu et al., 2014), the E/I ratio was similar for 1-and 2-minute NMDA application. This suggests that FS interneurons maintain a stable ratio of inhibitory and excitatory inputs despite excitatory and inhibitory plasticity.

It is particularly interesting that the patterns of coplasticity in *so*INs differ markedly from those in CA1 pyramidal cells. Indeed, while in pyramidal neurons, the coplasticity induced by 1-minute NMDA application preserves E/I balance (Wiera et al., 2024), this stimulation in the *stratum oriens* interneurons shifts the E/I balance in favor of excitation, likely promoting enhanced feedback and feedforward inhibition at the network level. However, following prolonged NMDA application, pyramidal cells exhibit asymmetric coplasticity with enhanced inhibition relative to excitation, which is in stark contrast to interneurons, for which iLTD is observed while excitatory inputs remain stable or slightly potentiated.

The observed plasticity dynamics in *so*INs suggests a potential network mechanism in which OLM/bistratified interneurons play a pivotal role in regulating the associativity of CA1 pyramidal cells. During brief NMDA receptor activation in CA1 INs, asymmetric coplasticity (eLTP and iLTD) in OLM/bistratified cells facilitates their disinhibition, allowing these interneurons to respond more robustly to excitatory inputs. Simultaneously, coplasticity in pyramidal cells upregulates both inhibitory and excitatory drives (Wiera et al. 2024). Thus, not only are OLM/bistratified cells disinhibited in response to NMDA, but this stimulus also upregulates the strength of their input to pyramidal neurons. Together, these mechanisms strengthen the feedback inhibitory control exerted by SST+ OLM/bistratified cells over the dendrites of pyramidal neurons. This enhanced inhibitory regulation may, in turn, modulate active Ca²⁺ events in pyramidal cell dendrites, which are essential for processes such as behavioral time scale plasticity and place cell formation (Bittner et al., 2017; Rolotti et al., 2022).

In conclusion, we have demonstrated the critical role of NMDA receptor activation in shaping both GABAergic synaptic plasticity at VIP-mediated inhibitory inputs and inhibitory-excitatory coplasticity measured at OLM/bistratified and FS interneurons in the CA1 *stratum oriens*. These processes are vital for modulating the excitation/inhibition balance, and understanding them may provide further insights into how coplasticity in various interneurons influences hippocampal network activity, place cell formation and associative learning. Future studies should focus on uncovering the intersynaptic molecular mechanisms that drive different coplasticity patterns and explore their role in network-level dynamics, particularly how I→I GABAergic plasticity contributes to disinhibitory processes relevant to behavior.

## Author contributions

Conceptualization: G.W and J.W.M. Investigation and data analysis: J.J., G.W., and J.W.M. Supervision: G.W. and J.W.M. Funding acquisition: J.W.M. Writing the draft and editing: J.J., G.W. and J.W.M.

## Acknowledgments

This work was supported by funding from the National Science Centre (Poland) grant OPUS 2021/43/B/NZ4/01675.

