## Supplement for "NMDA-dependent coplasticity in VIP interneuron-driven inhibitory circuits"

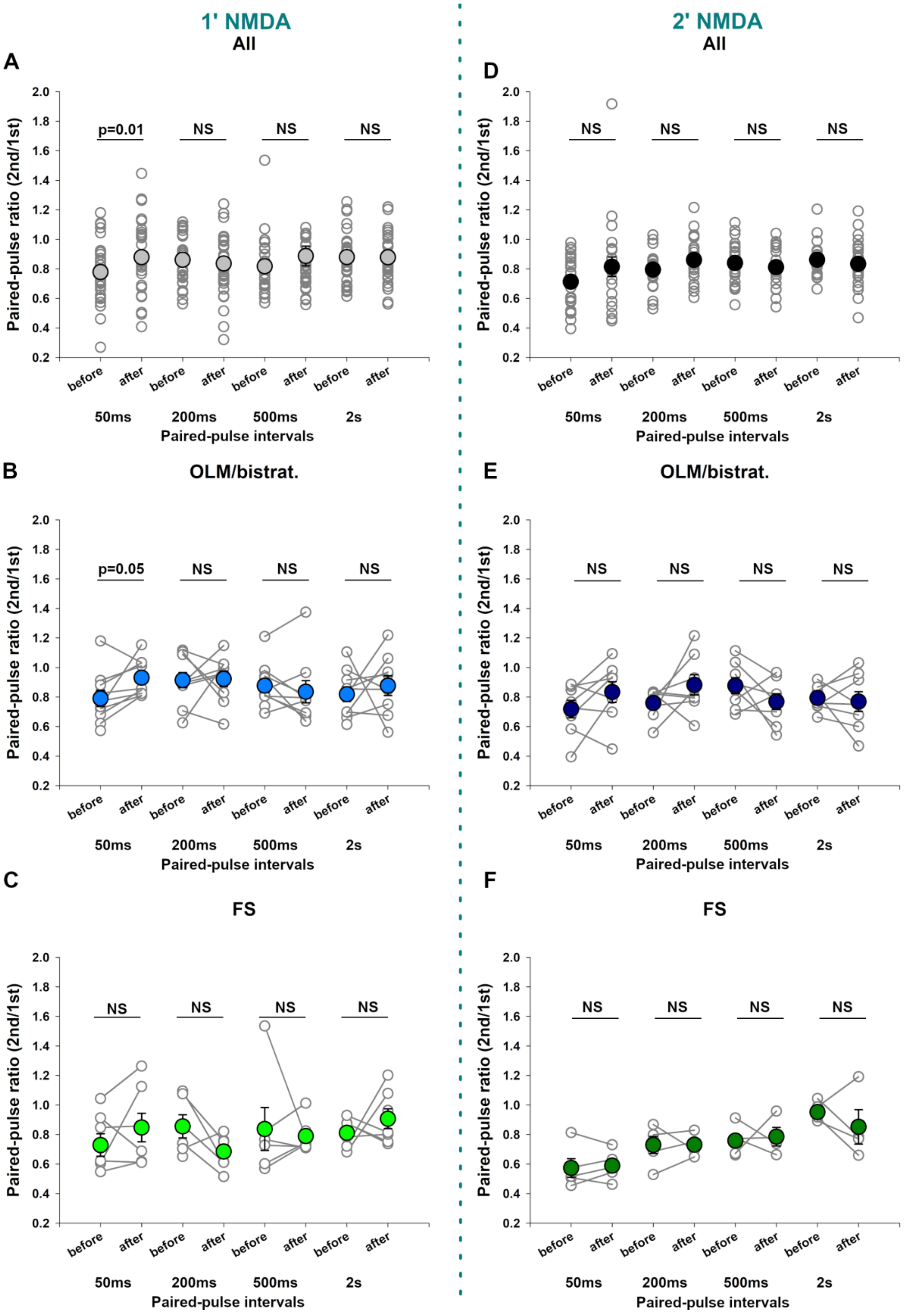


**Supplemental Figure 1. Comparison of VIP→*so*IN paired-pulse ratios before and after NMDA application.**

**(A-C)** Analysis of paired-pulse responses in VIP→*so*INs synapses measured before and 30 min after NMDA (1 min) administration. (**A**) Cumulative data for all recorded CA1 *stratum oriens* interneurons, regardless of interneuron class (50 ms before: 0.78 ± 0.03, after: 0.88 ± 0.04, $p$ = 0.01; 200 ms before: 0.86 ± 0.03, after: 0.84 ± 0.04, $p$ = 0.6; 500 ms before: 0.81 ± 0.03, after: 0.89 ± 0.07, $p$ = 0.35; 2 s before: 0.88 ± 0.03, after: 0.88 ± 0.03, $p$ = 0.8); (**B**) Comparison of paired-pulse ratios for OLM/bistratified cells (50 ms before: 0.79 ± 0.06, after: 0.93 ± 0.04, $p$ = 0.049; 200 ms before: 0.91 ± 0.05, after: 0.92 ± 0.05, $p$ = 0.9; 500 ms before: 0.88 ± 0.05, after: 0.83 ± 0.07, $p$ = 0.3; 2 s before: 0.82 ± 0.05, after: 0.88 ± 0.07, *p* = 0.71); (**C**) Comparison within the identified class of postsynaptic fast-spiking interneurons (50 ms before: 0.73 ± 0.08, after: 0.85 ± 0.1, *p* = 0.14; 200 ms before: 0.86 ± 0.08, after: 0.68 ± 0.04, *p* = 0.11; 500 ms before: 0.84 ± 0.14, after: 0.79 ± 0.04, *p* = 0.7; 2 s before: 0.81 ± 0.04, after: 0.91 ± 0.07, *p* = 0.48).

(**D-F**) Analysis of paired-pulse responses in VIP→*so*INs synapses measured before and 30 min after NMDA (2 min) administration. (**D**) Cumulative data for all recorded CA1 *stratum oriens* interneurons, regardless of interneuron class (50 ms before: 0.71 ± 0.04, after: 0.81 ± 0.07, *p* = 0.08; 200 ms before: 0.8 ± 0.03, after: 0.86 ± 0.03, *p* = 0.12; 500 ms before: 0.84 ± 0.03, after: 0.81 ± 0.03, *p* = 0.38; 2 s before: 0.86 ± 0.03, after: 0.83 ± 0.04, *p* = 0.52); (**E**) Comparison of paired-pulse ratios for OLM/bistratified cells (50 ms before: 0.72 ± 0.06, after: 0.83 ± 0.07, *p* = 0.14; 200 ms before: 0.76 ± 0.03, after: 0.88 ± 0.07, *p* = 0.19; 500 ms before: 0.88 ± 0.05, after: 0.77 ± 0.05, *p* = 0.18; 2 s before: 0.79 ± 0.03, after: 0.77 ± 0.07, *p* = 0.7): (**F**) Comparison within the identified class of postsynaptic fast-spiking interneurons (50 ms before: 0.55 ± 0.06, after: 0.59 ± 0.04, *p* = 0.66; 200 ms before: 0.73 ± 0.04, after: 0.73 ± 0.04, *p* = 0.87; 500 ms before: 0.76 ± 0.05, after: 0.79 ± 0.06, *p* = 0.98; 2 s before: 0.95 ± 0.03, after: 0.85 ± 0.12, *p* = 0.39); 1-minute NMDA treatment reduces paired-pulse-induced depression of VIP→OLM/bistratified IN IPSCs at the 50 ms interval, while paired-pulse responses with longer intervals remain unaffected. Extended NMDA exposure does not significantly alter the paired-pulse ratio. The significance levels are indicated in each plot (paired *t*-test).

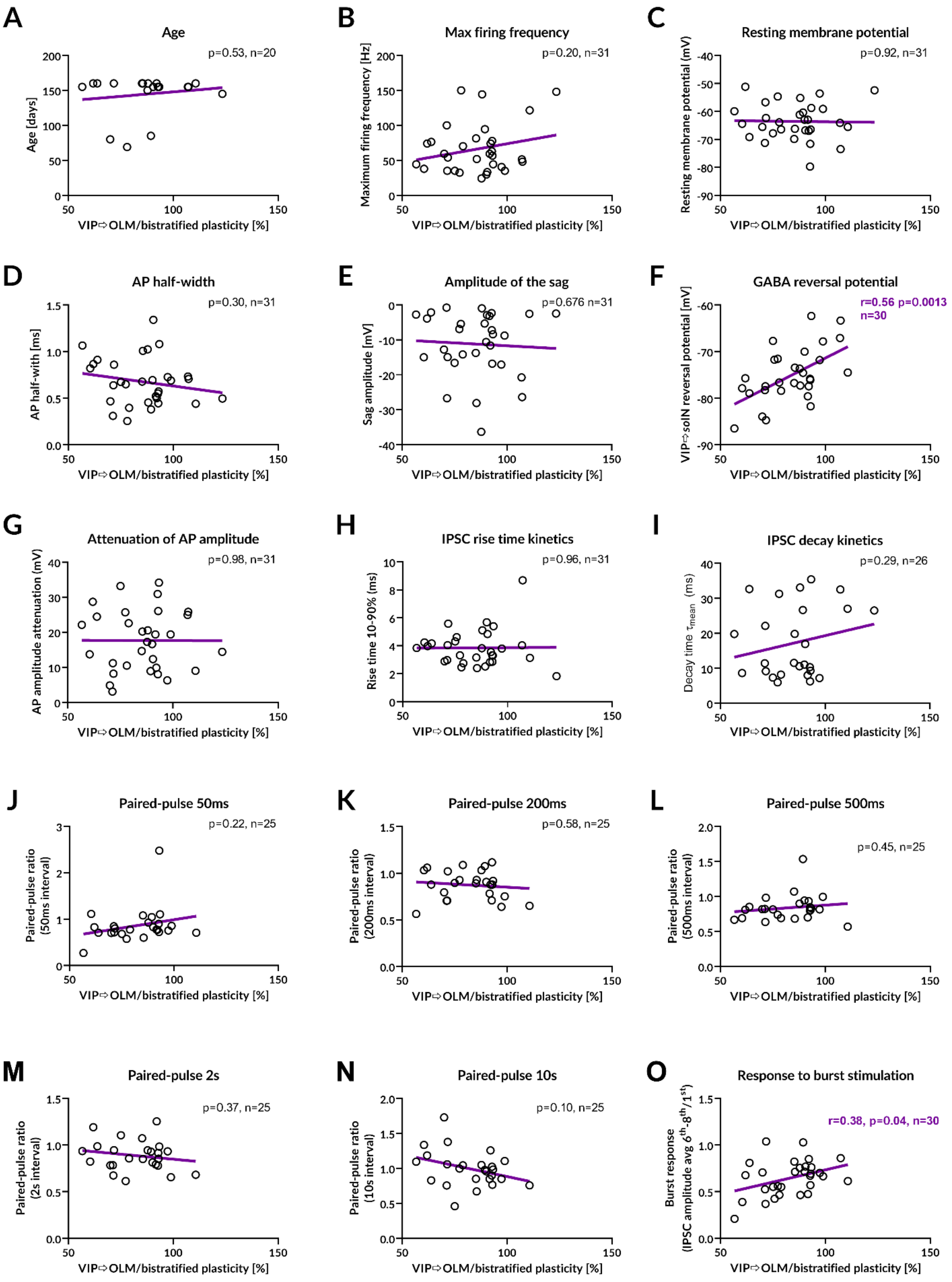


**Supplemental Figure 2. Correlation analysis of GABAergic plasticity in VIP inputs to the CA1 *stratum oriens* interneurons with various factors. Long-term plasticity was induced by 1-minute NMDA application.** The magnitude of plasticity was analyzed against (**A**) animal age, (**B-G**) electrophysiological properties of postsynaptic interneurons, (**F**) reversal potential of GABAergic synaptic currents, (**H-I**) IPSC kinetics, (**J-N**) short-term plasticity measured as paired-pulse ratios across different interstimulus intervals, and (**O**) burst-induced depression. All plots represent data from experiments in which NMDA was applied for 1 min, a protocol that reliably induced significant iLTD in the studied VIP projections. Each plot includes the corresponding p statistics and sample sizes (n) for correlation analysis. When a significant correlation was observed, the Pearson's correlation coefficient (r) is reported.

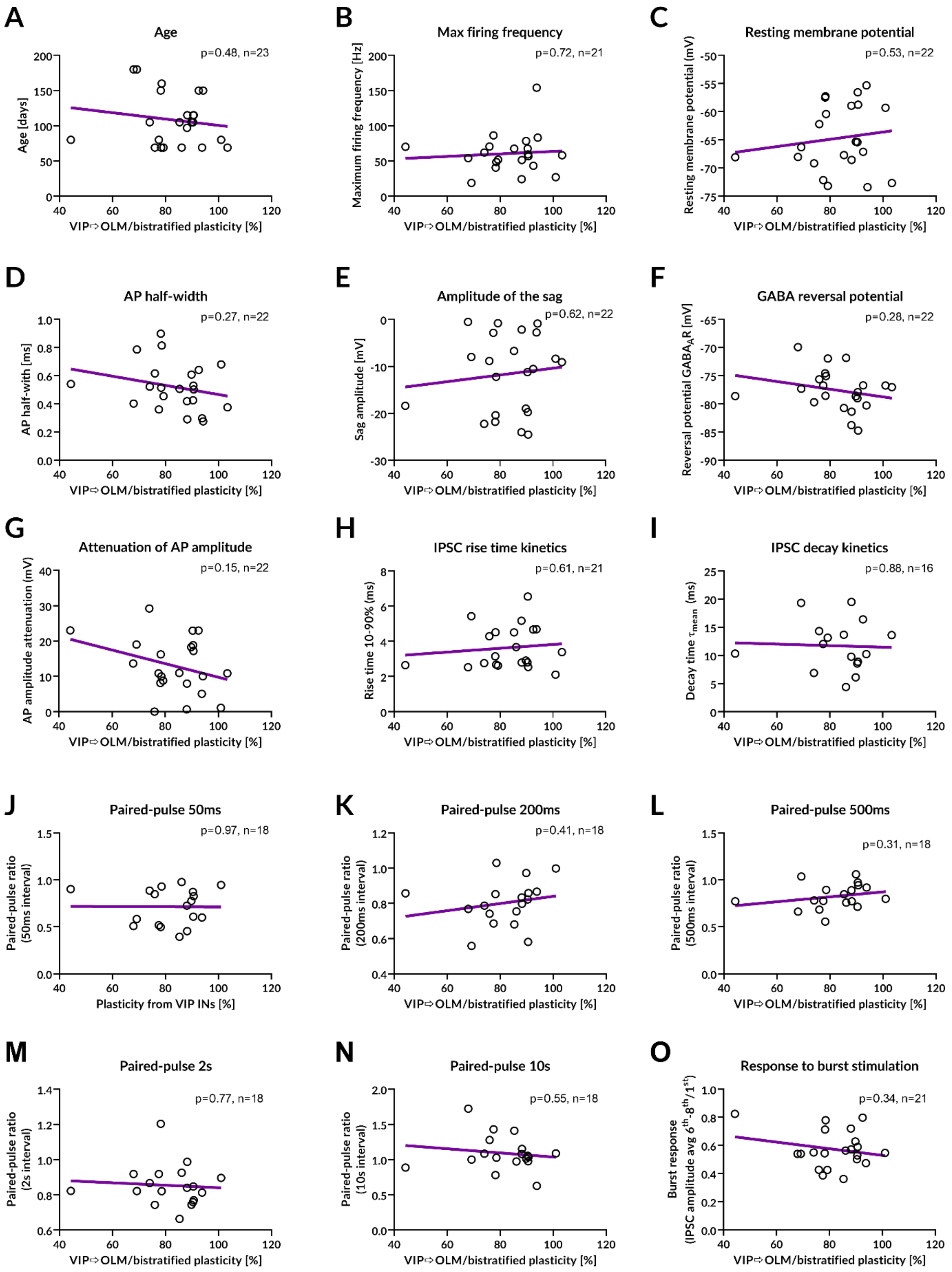


**Supplemental Figure 3. Correlation analysis of GABAergic plasticity in VIP inputs to the CA1 *stratum oriens* interneurons with various factors. Long-term plasticity was induced using a 2-minute NMDA application.** The magnitude of plasticity was analyzed in relation to (**A**) animal age, (**B-G**) electrophysiological properties of postsynaptic interneurons, (**F**) reversal potential of GABAergic synaptic currents, (**H-I**) IPSC kinetics, (**J-N**) short-term plasticity measured as paired-pulse ratios across different interstimulus intervals, and (**O**) burst-induced synaptic depression. All plots represent data from experiments using the NMDA application protocol, which reliably induced significant iLTD in the analyzed VIP projections. Each plot includes the corresponding p statistics and sample sizes (n) for correlation analysis. No significant correlations were observed.

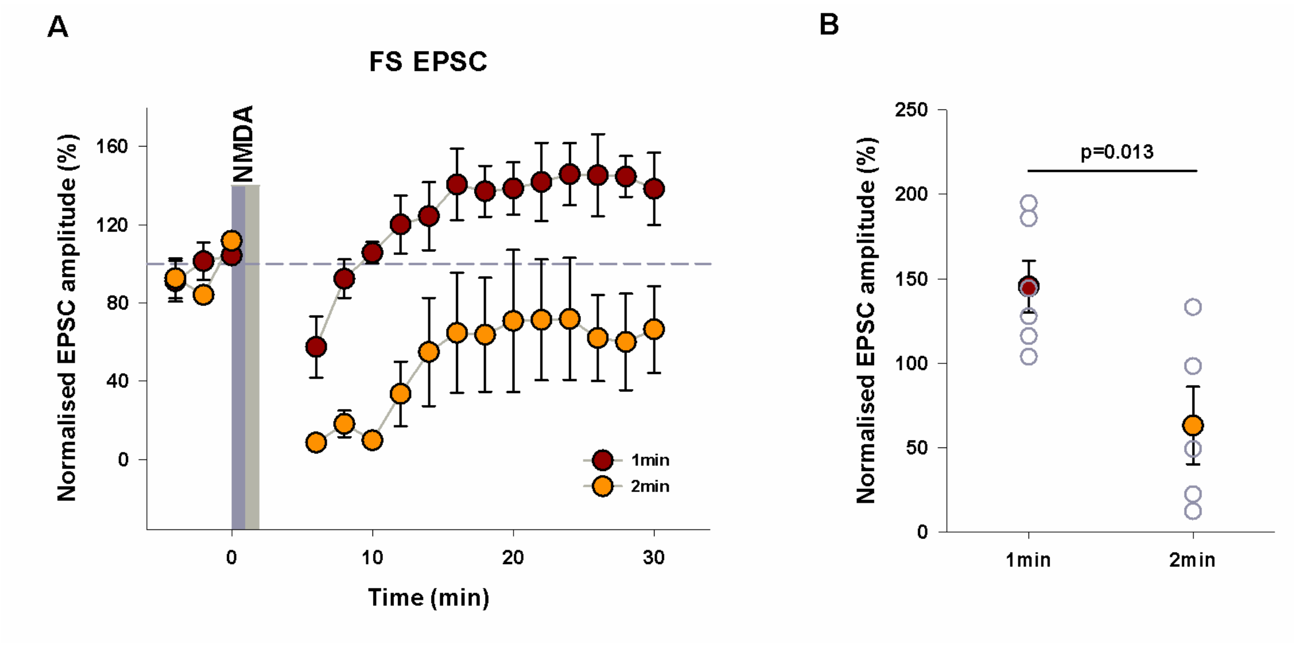


**Supplemental Figure 4. Comparison of NMDA-dependent long-term plasticity of excitatory inputs to fast-spiking interneurons in the CA1 *stratum oriens*.**

**(A)** Long-term plasticity at excitatory synapses onto *stratum oriens* fast-spiking (FS) interneurons, induced by 1-minute or 2-minute NMDA application. Excitatory synaptic transmission was evoked by placing a stimulation electrode in the CA1 stratum radiatum.

**(B)** Normalized percentage changes in EPSC amplitudes recorded from *stratum oriens* FS interneurons. A shorter (1-minute) NMDA application induced excitatory LTP, whereas extending the stimulation to 2 min resulted in a reversal of plasticity direction, leading to excitatory LTD. This figure includes the data presented in Figs. 2 and 3.
